# TinkerHap - A Novel Read-Based Phasing Algorithm with Integrated Multi-Method Support for Enhanced Accuracy

**DOI:** 10.1101/2025.02.16.638517

**Authors:** Uri Hartmann, Eran Shaham, Dafna Nathan, Ilana Blech, Danny Zeevi

**Affiliations:** Department of Biotechnology, Jerusalem Multidisciplinary College, Jerusalem, Israel

## Abstract

Phasing, the assignment of alleles to their respective parental chromosomes, is fundamental to studying genetic variation and identifying disease-causing variants. Traditional approaches, including statistical, pedigree-based, and read-based phasing, face challenges such as limited accuracy for rare variants, reliance on external reference panels, and constraints in regions with sparse genetic variation.

To address these limitations, we developed TinkerHap, a novel and unique phasing algorithm that integrates a read-based phaser, based on a pairwise distance-based unsupervised classification, with external phased data, such as statistical or pedigree phasing. We evaluated TinkerHap’s performance against other phasing algorithms using 1,040 parent-offspring trios from the UK Biobank (Illumina short-reads) and GIAB Ashkenazi trio (PacBio long-reads). TinkerHap’s read-based phaser alone achieved higher phasing accuracies than all other algorithms with 95.1% for short-reads (second best: 94.8%) and 97.5% for long-reads (second best: 95.5%). Its hybrid approach further enhanced short-read performance to 96.3% accuracy and was able to phase 99.5% of all heterozygous sites. TinkerHap also extended haplotype block sizes to a median of 79,449 base-pairs for long-reads (second best: 68,303 bp) and demonstrated higher accuracy for both SNPs and indels. This combination of a robust read-based algorithm and hybrid strategy makes TinkerHap a uniquely powerful tool for genomic analyses.

## Introduction

Phasing is the process of assigning alleles to their respective maternal or paternal chromosomes. It is essential for determining precise protein sequences in an individual and identifying genes that cause diseases.

Various methods of phasing are available, including statistical phasing based on phased reference genomes (e.g. ShapeIT [1] and Beagle [2]), pedigree-based phasing (e.g. LINKPHASE3 [3] and TrioPhaser [4]), and read-based phasing (e.g. WhatsHap [5] and HapCUT2 [6]). Statistical phasing is limited by how well the reference panel represents the sample data and is particularly inaccurate in phasing rare variants [7]. Pedigree-based phasing is accurate for common and rare variants, but pedigree information is typically not available for the inspected individual.

Read-based (or read-aware) phasing works by analyzing sequencing reads that span multiple heterozygous sites to phase them together, resulting in very high accuracy. This method is independent of reference bias, remains unaffected by the rarity of the alleles, and does not require pedigree data. However, since read-based phasing cannot phase regions where heterozygous sites are farther than read sizes, it can only effectively be used in highly variable regions or when using long-reads that span several heterozygous sites.

Here we present TinkerHap, a read-based phasing tool offering consistent performance and enhanced accuracy. TinkerHap excels in accurately handling rare variants and variable genomic regions, such as the Human Leukocyte Antigen (HLA) locus, while also effectively phasing long-read data. Moreover, TinkerHap uniquely integrates information from statistical phasing methods into its read-based framework. This hybrid approach bridges gaps in read coverage and extends haplotype blocks.

## Methods

### Overview

TinkerHap is implemented in Python 3 and utilizes the *pysam* package [8] for manipulating alignment and variant calling files. The command-line interface accepts an alignment file (SAM/BAM/CRAM) and a variant calling file (VCF/BCF) as inputs, producing a phased VCF file through read-based haplotype phasing. Optionally, TinkerHap can integrate a pre-phased VCF file from a third-party tool (e.g., ShapeIT [1] for statistical-based phasing) to align and merge haplotypes with greater accuracy when possible.

Additionally, TinkerHap can generate multiple output formats to represent the phased haplotypes: a BED file listing the identified haplotype blocks, a BAM file identical to the original but annotated with haplotype and phase information in the Haplotype Phase field (HP) and Haplotype number field (HT), and two separate BAM files - each containing reads corresponding to one of the phased alleles. These outputs facilitate annotation or the splitting of the original alignment into distinct files for each allele, enabling downstream analyses.

### Algorithm

TinkerHap implements a three-step phasing algorithm, based on a pairwise distance-based unsupervised classification, designed for precision and scalability. Below is a detailed description of each step with mathematical notations.

#### 1. Identification of Heterozygous Sites

Let *S* = {*s*_1_, *s*_2_,…, *s*_*m*_} represent the set of heterozygous sites identified from the input variant call file (VCF). A site *s*_*i*_ is considered heterozygous if *a*_*i*_ ≠ *b*_*i*_ where *a*_*i*_ and *b*_*i*_ are the two alleles at *s*_*i*_. The loci of these sites are identified as *L* (*S*) = {*l*_1_, *l*_2_,…, *l*_*m*_}, forming the foundation for subsequent phasing steps. Ambiguous allele calls, characterized by low scores in the VCF’s QUAL column and typically caused by sequencing or alignment errors, are flagged.

#### 2. Association of Reads with Heterozygous Sites

Let *R* = {*r*_1_, *r*_2_,…, *r*_*m*_} denote the set of sequencing reads. Each read *r*_*j*_ spans a subset of heterozygous sites *S*_*j*_ ⊆ *S*. For each read, we map its alleles to the overlapping sites:

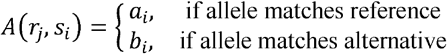

This step ensures precise allele identification by linking reads to heterozygous sites while accounting for potential alignment errors or ambiguities, such as indels. To ensure accuracy, only reads meeting a minimum mapping quality threshold (e.g., MAPQ ≥ 20) are considered, ensuring that low-confidence alignments do not influence the phasing process.

#### 3. Calculation of Phase Scores

The phasing process begins by arbitrarily assigning the first read *r*_1_ to one of the haplotypes, for instance *H*_1_. This initial assignment acts as a seed to propagate haplotypes across all overlapping reads.

Each read *r* is then evaluated to determine its phase matching scores *P*_1_(*r*) and *P*_2_(*r*) for the two haplotypes, *H*_1_ and *H*_2_. These scores are computed by analyzing all overlapping reads and the heterozygous sites they share with *r*.

The phase scores are calculated as:

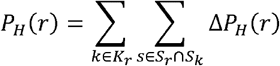

Where *K*_*r*_ is the set of all overlapping reads for *r*, and *S*_*r*_ and *S*_*k*_ are the sets of heterozygous sites for reads *r* and *k*, respectively. The contribution of each shared site *s* to the phase score, Δ*P*_*H*_(*r*), is defined as:

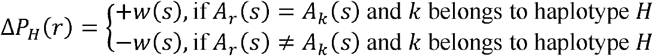

Here, *A*_*r*_(*s*) and *A*_*k*_(*s*) represent the alleles of *r* and *k* at site *s*, respectively. The weight *w*(*s*) assigned to site *s* depends on the type of heterozygous site:

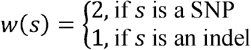

This scoring ensures that the phase scores for *r* are influenced by the agreement or disagreement between *r* and all overlapping reads at shared heterozygous sites.

After calculating the phase scores, the haplotype *HP* of *r* is assigned as follows:

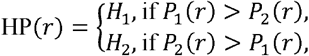

If *P*_1_(*r*) and *P*_2_(*r*) are equal, the haplotype assignment can propagate from the overlapping read with the strongest phase connection, or a new haplotype block may be started. This approach ensures consistency in haplotype assignments based on the majority consensus among overlapping reads.

#### 4. Haplotype Extension

Haplotypes are extended iteratively by analyzing overlapping reads. If a read *r*_*k*_ overlaps two or more phased reads 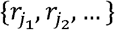, its phase is determined by propagating the majority consensus:

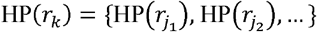

This ensures the consistency of haplotype assignments across contiguous genomic regions. Reads that span conflicting haplotypes are flagged for manual review or downstream quality filtering.

#### 5. Pair-End Read Merging

For paired-end reads (*r*_*j*,_ *r*_*j*_,) the algorithm evaluates the consistency of their haplotypes:

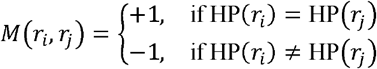

Inconsistent pairs trigger a phase reassignment to minimize discordance, leveraging the paired-end linkage information. A weighted graph representation of pair-end links can be constructed for further optimization of haplotype continuity.

#### 6. Integration with Pre-Phased Data (Optional)

When an additional pre-phased VCF file is provided, for example one generated by statistical phasing tools such as ShapeIT, the algorithm merges the read-based haplotypes with the pre-phased data. This involves merging haplotypes and, if necessary, switching the phase numbers (e.g., swapping haplotype 1 and haplotype 2) to ensure consistency with the pre-phased data numbering of haplotypes.

The alignment score *A* (*b,h*) for a pre-phased block *b* and a read-based block *h* is calculated as:

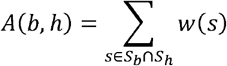

where *S*_*b*_ and *S*_*h*_ are the sets of heterozygous sites in *b* and *h*, and *w*(*s*) represents the weight based on the site type (e.g., SNP or indel). Haplotypes are adjusted to maximize *A* (*b,h*), ensuring that the merged haplotypes align with the pre-phased data and improving the overall phasing accuracy.

#### 7. Output Generation

The final outputs include:

1. Phased VCF: Annotated with a PS (Phase Set) field.
2. Annotated BAM: Each read is tagged with HP (Haplotype Phase) and HT (Haplotype number) fields.
3. Split BAM Files: Separate BAM files for each haplotype, facilitating downstream analyses.
4. BED File: Haplotype boundaries across the genome are defined for visualization.

The detailed algorithm and code can be accessed at: https://github.com/DZeevi-Lab/TinkerHap

### Evaluation

TinkerHap was evaluated in the following use cases:

1. To evaluate the algorithm’s performance in variable regions and for rare variants using Illumina short-reads, we analyzed Whole Genome Sequencing (WGS) data from 1,040 parent-offspring trios that we identified in the UK Biobank [9] on the MHC class II region in humans, specifically on chr6:32,439,878-33,143,325 (hg38 genome version).
2. To evaluate the algorithm’s performance with long-reads, we used PacBio long-read sequencing data of the full genomes of GIAB Ashkenazi trio HG002-4 and Chinese trio HG005-7 datasets by Revio (publicly offered by GIAB [10]).

For each offspring in the trios, we constructed a “truth” set of known phased heterozygous sites (“truth sites”). This was achieved by examining loci where each parent possesses different homozygous alleles or where one parent was heterozygous, and the other was homozygous. After preparing the data, we phased the offspring sequence using the following algorithms: ShapeIT[1], WhatsHap[5], HapCUT2[6], TinkerHap, and TinkerHap with ShapeIT[1] phased data as an additional input for merging haplotypes (as described in the “Algorithm” section above). The success rate was evaluated by counting the number of sites in the phased output that matched the truth set.

## Results

### MHC class II gene region phasing (chr6:32,439,878-33,143,325)

**Table 1.** Phasing performance of different algorithms on short-reads aligned to the MHC class II region. ^1^ TinkerHap+ShapeIT: TinkerHap algorithm when used with additional ShapeIT pre-phased file. ^2^ Phasing accuracy: Successfully phased sites divided by the total number of heterozygous sites. ^3^ Haplotype size: Median haplotype size across all samples. “Haplotype” refers to a set of alleles at variant sites along a single chromosome that are inherited together and are guaranteed to be phased together by the algorithm. ^4^ Common phased sites errors: Phasing error % in heterozygous sites phased by all algorithms. ^5^ Runtime: Median runtime per sample.

| Criteria | TinkerHap+<br>ShapeIT<sup>1</sup> | TinkerHap | WhatsHap | HapCUT2 | ShapeIT |
| --- | --- | --- | --- | --- | --- |
| **Phased %** | 99.5% | 97.1% | 86.5% | 96.2% | 70.5% |
| **Phasing accuracy %<sup>2</sup>** | 96.3% | 95.1% | 84.9% | 94.8% | 70.2% |
| **Phasing accuracy % (SNPs only)** | 97.1% | 96.0% | 86.0% | 95.8% | 71.6% |
| **Phasing accuracy % (INDELs only)** | 89.6% | 87.8% | 76.9% | 87.4% | 59.6% |
| **Haplotype size (bp)<sup>3</sup>** | 21,813 | 631 | 75 | 644 | 702,123 |
| **Common phased sites errors<sup>4</sup>** | 0.14% | 0.14% | 0.18% | 0.12% | 0.17% |
| **Runtime (s)<sup>5</sup>** | 6.8 | 6.5 | 23.8 | 9.2 | 13.7 |
| **Bases / second** | 104,005 | 108,245 | 29,532 | 76,105 | 51,545 |
| **Heterozygous sites / second** | 743 | 773 | 211 | 544 | 368 |
| **Memory usage (MB)** | 109 | 106 | 397 | 594 | 24 |

**Figure 1.**
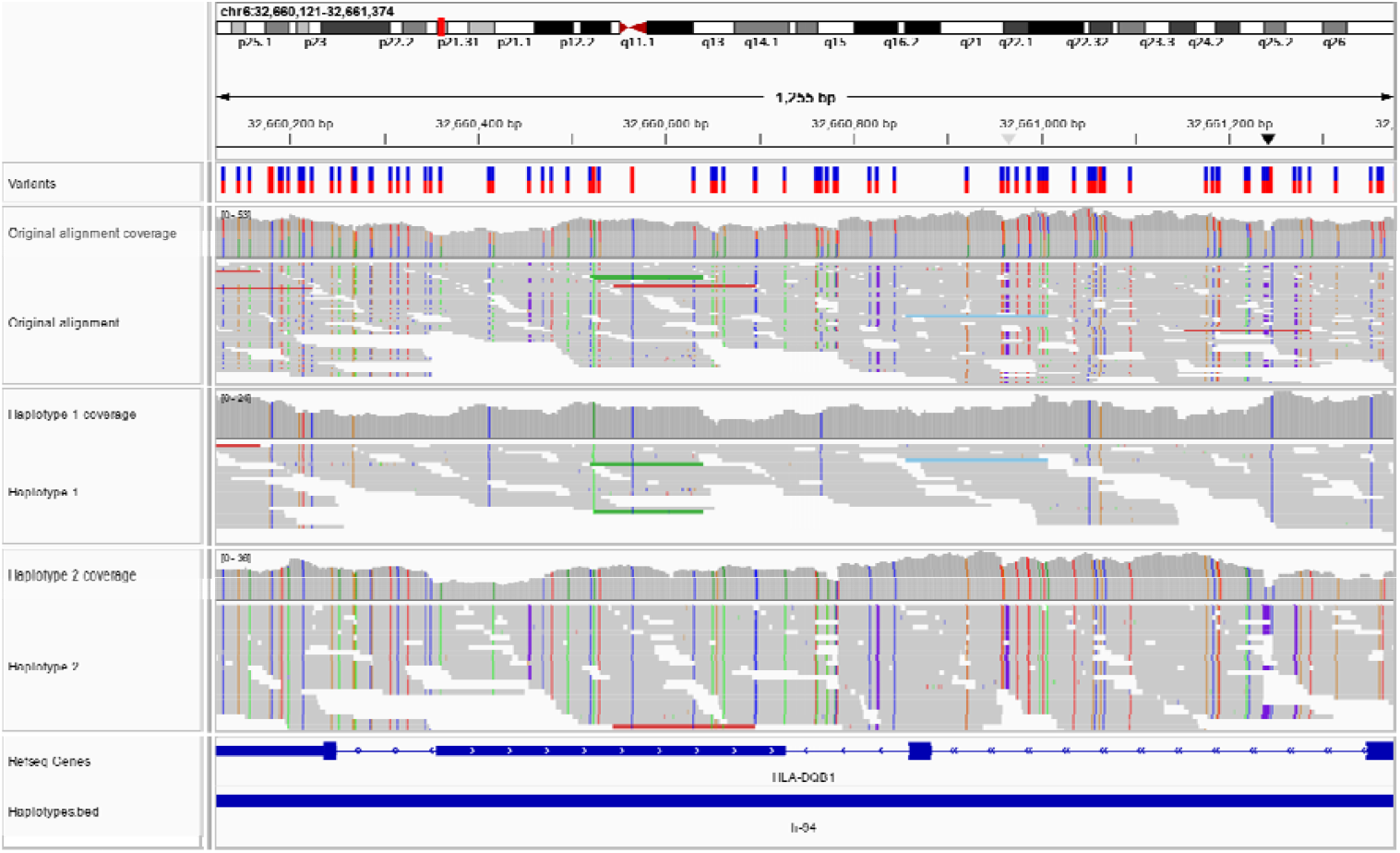
Phased BAM Outputs displaying Heterozygous sites. (IGV [11] screenshot) - The upper track is the original alignment, while the two tracks below represent the output of Haplotype 1 and Haplotype 2. Heterozygous sites are correctly segregated between the two phases, demonstrating successful phasing. The continuous blue line in the bottom track illustrates a BED file annotation, highlighting the size of the haplotype region where all variants are confirmed to share the same phase.

### PacBio phasing (whole genome)

**Table 2.** Phasing performance of different algorithms on long-reads of full genomes. ^1^ Phasing accuracy: Successfully phased sites divided by the total number of heterozygous sites. ^2^ Haplotype size: Median haplotype size across all samples. ^3^ Common phased sites errors: Phasing error % in heterozygous sites phased by all algorithms. ^4^ Runtime: Median runtime per sample.

| Criteria | TinkerHap | WhatsHap | HapCUT2 |
| --- | --- | --- | --- |
| **Phased %** | 99.4% | 96.8% | 96.7% |
| **Phasing accuracy %<sup>1</sup>** | 97.5% | 95.5% | 95.4% |
| **Phasing accuracy % (SNPs only)** | 97.8% | 96.5% | 96.4% |
| **Phasing accuracy % (INDELs only)** | 96.0% | 90.2% | 90.5% |
| **Haplotype size (bp)<sup>2</sup>** | 79,449 | 68,303 | 72,220 |
| **Common phased sites errors<sup>3</sup>** | 0.47% | 0.47% | 0.55% |
| **Runtime (s)<sup>4</sup>** | 10,519 | 17,578 | 5,495 |
| **Bases / second** | 298,239 | 178,475 | 570,891 |
| **Heterozygous sites / second** | 433 | 259 | 829 |
| **Memory usage (MB)** | 1,726 | 1,696 | 876 |

## Discussion

Here, we introduce TinkerHap, a read-based phasing algorithm designed for accurate and reliable phasing across diverse genomic contexts, with the ability to integrate statistical phasing data from third-party tools for improved performance.

We evaluated TinkerHap using two datasets: the MHC class II region in humans with Illumina short-read WGS data to assess its accuracy in variable regions, and PacBio sequencing data to evaluate its performance with long-reads. These datasets were selected due to their suitability for testing read-based phasing algorithms, as both are characterized by a high density of variants that provide many opportunities for phasing.

### Performance of Short-Reads in Variable Regions

In the MHC class II region using short-reads, TinkerHap phased 97.1% of variants with 95.1% accuracy. In comparison, the second-best algorithm phased 96.2% of variants with 94.8% accuracy. All methods showed higher phasing accuracy for SNPs compared to indels (97.1% and 89.6%, respectively, in TinkerHap).

### Performance of Long-Read Sequencing

TinkerHap achieved a phasing accuracy of 97.5% for SNPs and 96.0% for indels with PacBio datasets. These results were superior to the second-best algorithm, which demonstrated accuracies of 95.5% and 95.4%, respectively. Moreover, TinkerHap produced longer haplotype blocks (median size: 79,449 bp) compared to the second-best algorithm (68,303 bp). Runtime analysis revealed that TinkerHap required 10,519 seconds per sample, compared to 5,495 seconds for the fastest algorithm.

### Comparison of Long-Read and Short-Read Performance

TinkerHap performed better with long-read sequencing data compared to short-read data in several key metrics. Long-reads offer superior upstream alignment quality, particularly at highly variable sites, which enhances the overall accuracy of variant calling and subsequent phasing steps. Long-read data yielded more extensive haplotype blocks (median size: 79,449 bp compared to 631 bp with short-reads) and higher phasing accuracy (97.5% for SNPs in long-reads compared to 96.0% in short-reads, and 96% for indels in long-reads compared to 87.8% in short-reads). This improved performance is expected due to long-reads containing more heterozygote sites and enabling improved alignments.

### Integration with Statistical Phasing

TinkerHap uniquely includes the ability to integrate data from third-party tools, such as ShapeIT. By incorporating pre-phased haplotypes, the TinkerHap + ShapeIT combination achieved 99.5% phased variants with 96.3% accuracy, significantly outperforming standalone methods. This hybrid approach improved haplotype block continuity and effectively addressed gaps in read coverage.

### Limitations

TinkerHap’s runtime and memory usage for long-read data present areas for potential optimization, and it currently lacks support for polyploid genomes. TinkerHap is limited in merging distant haplotypes, which could be particularly useful for applications such as exome sequencing. Future incorporation of pedigree information could address this issue and enhance TinkerHap’s accuracy in trio or family-based studies.

In most phasing errors that we manually examined, inaccuracies were primarily attributed to upstream variant calling rather than to the phasing algorithm itself. This suggests that TinkerHap may be approaching the limit of what can be achieved with downstream read-based phasing alone. This underscores the importance of high-quality preprocessing.

## Author contribution

Conceptualization – U.H., D.Z.; Methodology - U.H.; Formal Analysis - U.H.; Investigation - U.H; Writing, original draft preparation – U.H.; Writing, review & editing - U.H., D.Z., E.S., D.N, I.B.; Visualization - U.H. Supervision – D.Z.; Funding Acquisition – D.Z.

## Acknowledgements

We thank Rona Gershon Talmi from the Hamaabada Podcast (Kan) and Dr. Jeremy Fogel and Tuval Rosenwasser from the Think & Drink Different Podcast for their contribution to this work.

This research has been conducted using the UK Biobank Resource under application number 74655.

